# Effects of lambda-cyhalothrin on gut microbiota and related bile acid metabolism in mice

**DOI:** 10.1101/2024.08.24.609540

**Authors:** Weijia Zheng, Lingyuan Xu, Maojun Jin, Jing Wang, Ivonne M.C.M. Rietjens

## Abstract

**Background:** Since the gut microbiota plays a crucial role in host metabolism and homeostasis, its alterations induced by xenobiotics, such as pesticides, could pose a risk to host health. Our previous in vitro fermentation study showed that pyrethroid pesticides could affect the mouse bacterial community and related bile acid profiles. Hence in the present study, the effects of the selected pyrethroid lambda-cyhalothrin on the intestinal microbial community and its related bile acid metabolism were evaluated in male and female mice.

**Results:** The total amount of bile acids in plasma and fecal samples from lambda-cyhalothrin treated mice markedly increased compared to controls, which could be mainly ascribed to the significantly raised proportions of taurine conjugated bile acids in plasma, and the increase in fecal secondary bile acids. In gut microbial profiles, a significantly increased richness of *Prevotellacea* and a depletion of *Lachnospiraceae* were found at the family level upon the treatment with lambda-cyhalothrin.

**Conclusion:** Treatment of mice with lambda-cyhalothrin affected the gut microbiota with accompanying changes in bile acid homeostasis. The effects on fecal bile acid profiles were in line with those previously observed in our in vitro study and corroborate that pyrethroid pesticides could affect gut microbiota and related bile acid profiles.

## Introduction

Bile acids, which are synthesized from cholesterol in hepatocytes, secreted into bile, stored in the gallbladder of human, and released into the intestine via the bile duct, are essential for intestinal solubilization, absorption, and metabolism of triglycerides, cholesterol, and fat-soluble vitamins (Fiorucci & Distrutti, 2015; Jia et al, 2018). Bile acids produced in the liver are referred to as primary bile acids, and may differ between species, with cholic acid (CA) as well as ursodeoxycholic acid (CDCA) being produced in humans, and CA, CDCA, α-muricholate (αMCA) as well as β-muricholate (βMCA) in rodents (Sayin et al., 2013). Further, these primary bile acids are conjugated to either mainly glycine in humans or taurine in rodents (Liu et al, 2015).

Upon release into the intestine, these conjugated bile acids are deconjugated by bacterial enzymes providing bile salt hydrolase (BSH) activity, and further metabolized to secondary bile acids by dehydroxylation, hydroxylation, or epimerization (Martin et al., 2007), resulting in formation of for example DCA and LCA by dehydroxylation of respectively CA and CDCA in humans or of ωMCA by 6β-epimerization of βMCA in rodents (Thomas et al, 2008). Despite trillions of bacteria in the gut, only a few species have been reported to have the capacity for transforming bile acids, such as *Clostridium* and *Bacteroides* which are able to support deconjugation by BSH as well as epimerization (Jia et al, 2018). Given this capacity of the gut microbiota for bile acid metabolism, the composition of the gut microbiota is a critical determinant in the ultimate bile acid profiles and related health aspects in the host (Fiorucci and Distrutti, 2015). It also implies that bile acid profiles may be disrupted by factors that affect the intestinal environment and thereby the gut microbiota (De Vadder et al., 2014).

Pyrethroids are extensively used pesticides which have been applied in agricultural and home formulations for three decades and account for approximately one-fourth of the worldwide insecticide market (Casida and Quistad, 1998). Pyrethroids can be classified into two large groups depending on their chemical structures without or with a cyano moiety (type I or type II), with cyhalothrin (**Fig.1**) being a classic type II pyrethroid. Cyhalothrin is a racemic mixture consisting of two pairs of enantiomeric isomers (**Fig. 1**) with pair B being lambda-cyhalothrin (Birolli et al., 2018), which is the active component of cyhalothrin and commonly used. The Joint Meeting on Pesticides Residues has set the ARfD of 0.02 mg /kg bw for lambda-cyhalothrin (JMPR 2018). The adverse health effects caused by these type II pyrethroids include neurotoxicity resulting in hypersensitivity, profuse salivation, choreoathetosis, tremor and paralysis (Vijverberg and vanden Bercken, 1990).

**Fig. 1.**
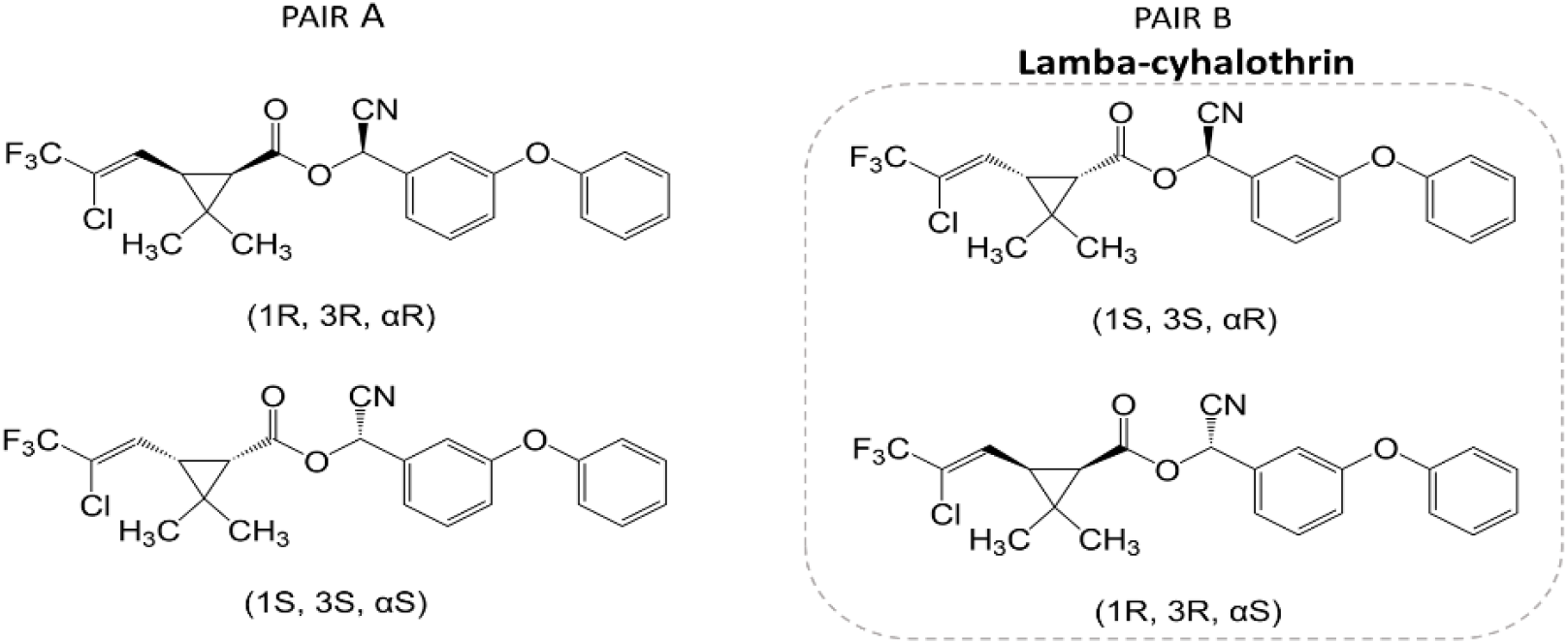
Chemical structure of the cyhalothrin stereoisomers including lambda-cyhalothrin (pair B).

The potential toxicity of pesticides towards the intestinal microbial community only started to gain attention in recent years (Giambò et al, 2021). Thus, studies published so far on gut microbiota toxicity of pyrethroids are limited. A study of Nasuti et al. (Nasuti et al., 2016) reported a decrease in the richness of the phyla of *Bacteroidota* in rat gut upon the administration of permethrin (a type I pyrethroid). Recently, another study reported that the exposure of small yellow croaker embryos to lambda-cyhalothrin markedly altered eight common genera of their gut microbial community, accompanied by significantly increased levels of the bile acids taurodeoxycholic acid (TDCA) and tauroursodeoxycholic acid (TUDCA) in water samples from their living surroundings (Zhan et al., 2022).

Moreover, in our previous work which utilized an optimized batch fermentation model system using mice feces, effects of the pyrethroids cypermethrin and cyhalothrin on gut microbiota-mediated bile acid profiles were assessed. In that study all pesticides appeared to affect the fecal gut microbiota and related bile acid metabolism, with the exposure to cyhalothrin resulting in the most significant alteration of fecal bile acid profiles. The exposure of cyhalothrin caused a significant increase in fecal secondary bile acids and an accompanying decrease in fecal primary bile acids, resulting in a raised ratio of βMCA to ωMCA. Results obtained from that in vitro fermentation batch model indicated that pesticides may affect intestinal microbiota and the related bile aid profiles. Hence the aim of the present study was to determine whether also in vivo such effects would occur. To that end C57BL/6 mice were exposed to lambda-cyhalothrin at two selected levels for a 28-day oral dose toxicity study to evaluate its influences on the intestinal microbial community, as well as on fecal and plasma bile acid profiles.

## Materials and methods

### Chemicals

Lambda-cyhalothrin was purchased from Sigma-Aldrich (St. Louis, MO, USA; CAS: 91465-08-6). Bile acid standards were supplied by Merck KGaA (Darmstadt, Germany), or Cambridge Isotope Laboratories (Tewksbury, USA). Formic acid and HPLC-grade acetonitrile were purchased from Merck (Darmstadt, Germany). Ultrapure water was acquired from a Milli-Q water system (Millipore, Billerica, MA, USA).

### Animals and exposure

Eight-week-old male and female C57BL/6 mice were purchased from the Vital River Laboratory Animal Technology Co., Ltd. (Beijing, China). All animal studies were performed in accordance with the Regulations for Care and Use of Laboratory Animals and Guideline for Ethical Review of Animals (China, GB/T 35892-2018) and the overall project protocols were approved by the Animal Ethics Committee of Beijing Institute of Technology. The accreditation number is SCXK-BIT-20210715010 promulgated by the Animal Ethics Committee of Beijing Institute of Technology. All animals were submitted to controlled temperature conditions (22–26 °C), humidity (50–60%), and light (12 h light/ 12 h dark) with the access to water and food ad libitum.

Dose levels equal to 1/20 and 1/10 of the reported LD50 of 19.9 mg/kg bw in mice (JMPR, 2007) for lambda-cyhalothrin were selected as the low- and high-oral doses for lambda-cyhalothrin respectively, resulting in dose levels of 1 and 2 mg/kg bw/day. Lambda-cyhalothrin was dissolved in dimethyl sulfoxide (DMSO) and further diluted in ultra-pure water to final concentrations of 1 and 2 mg/mL for dosing female mice, and 1.1 and 2.2 mg/mL for dosing males. Mice were divided over the different study groups based on similar weight distribution over the groups. The low dose of 1 mg/kg bw/day (named LCT group), the high dose of 2 mg/kg bw/day (named 2LCT group) or vehicle (control group) were administered by oral gavage (0.02 mL prepared lambda-cyhalothrin solution) once a day for 28 days (n=5 mice per sex per group).

### Clinical examinations

The mice were observed and checked daily for clinically abnormal signs and mortalities. On study day 6, 13 and 27, body weight as well as food consumption were determined before the daily oral gavage. At the end of the 28-day treatment, all mice were sacrificed by decapitation under isoflurane anesthesia.

### Sampling of blood and feces for bile acid profiling

On study day 7, 14 and 28 (morning 8:30 to 10:30 h), blood samples were taken from the tail vein of all mice (1.0 mL K-EDTA blood). The collected blood samples were centrifuged (12,000 g, 5 min, 4 °C), and the separated EDTA plasma samples (upper layers) were collected. Subsequently, 10 μL of each EDTA plasma sample was extracted with 400 μL acetonitrile, vortexed, and centrifuged (12,000 g, 5 min, 4 °C). Supernatants were collected in Eppendorf tubes, covered with nitrogen and frozen at −80 °C until bile acid analysis was performed.

On study day 28, after a fasting period of 16–20 h and one day after the last administration of lambda-cyhalothrin, feces were carefully removed during necropsy from the rectum of male and female mice. Fecal samples collected from individual mice were weighted, crushed, and extracted with acetonitrile (2 mL/ 1 mg feces) in an ultrasonic bath for 10 min following a 5 min centrifugation (21,5000 g, 4 °C). The supernatants were removed and stored at −80 °C until bile acid analysis was performed.

### Bile acid measurement in plasma and feces

Bile acid measurement was performed on a Nexera XR LC-20AD SR UPLC system coupled to a triple quadrupole liquid chromatography mass spectrometry (LCMS) 8050 mass spectrometer (Kyoto, Japan) with electrospray ionization (ESI) interface, which was able to measure the following 24 bile acids: 15 conjugated bile acids including T/G-CA, T/G-CDCA, T/G-DCA, T/G-LCA, T/G-UDCA, T/G-HDCA, TαMCA, TβMCA, and TωMCA; 4 primary bile acids including CA, CDCA, αMCA, βMCA; 5 secondary bile acids including ωMCA, DCA, LCA, UDCA, and HDCA. Prior to analysis all the extracts of plasma and fecal samples were shaken briefly, centrifuged (21,5000 g, 4 °C), and filtered through a 0.2 μm syringe filter (MILLEX, Merck Millipore, Cork, Ireland).

Bile acids in all samples were separated on a Kinetex C18 column (1.7μm×100 A×50 mm×2.1mm; Phenomenex, USA) using an ultra-high performance liquid chromatography (UHPLC) system (Shimadzu) with mobile phases consisting of 0.01% formic acid in distilled water (solvent A), a mixture of methanol and acetonitrile (v/v=1/1) (solvent B), and acetonitrile containing 0.1% formic acid (solvent C). The mass spectrometer (MS) used ESI in negative ion mode, and the parameters of ESI^-^ as well as the gradient profile are summarized in **Table S1**. Selective ion monitoring (SIM) and multiple reaction monitoring (MRM) were simultaneously employed for detection of the bile acids. The optimized precursor and product ions used for the detection of the bile acids are shown in **Table S1** as well. The Postrun and Browser Analysis function from the LabSolutions software (Shimadzu, Kyoto, Japan) was employed to obtain the peak areas of the extracted ion chromatogram (EIC) for each target.

### Bacterial profiling of gut microbiome

Microbial DNA was extracted from fecal samples of C57BL/6 mice using the QIAamp DNA Fecal Microkit (Stras Qiagen GMBH, Germany). A NanoDrop 2000 spectrophotometer (Thermoelectric Science, Massachusetts, USA) and 1 % agarose gel electrophoresis (AGE) was applied to measure the total DNA mass. Employing the extracted DNA as a template, an upstream primer 338F (5’-ACTCCTACGGGAGGCAGCAGCAG-3’) and a downstream primer 806R (5’ - GGACTACHVGGGTWTCTAAT-3’) with Barcode sequence were used to amplify the V3-V4 variable region of 16S rRNA genes (95 ºC reaction for 3 min, one cycle; denaturation at 95 ºC for 30 s, annealing at 55 ºC for 30 s, extension at 72 ºC for 45 s, 27 cycles; stably extended at 72 ºC for 10 min, ending at 10 ºC). The recovered products were purified using the AxyPrep DNA Gel Extraction Kit (Axygen Biosciences, Union City, CA, USA) and detected by 2 % AGE. The recovered products were detected and quantified using a Quantus™ Fluorometer (Promega, USA).

Purified amplicons were pooled in equimolar amounts and paired end sequenced on an Illumina MiSeq PE300 platform (Illumina, San Diego, USA) according to the standard protocols by Majorbio Bio-Pharm Technology Co. Ltd. (Shanghai, China). The data were then optimized by noise reduction processing to obtain ASV (Amplicon Sequence Variant) data representing the sequence and abundance information. The taxonomic identifications of 16S rRNA sequences were assigned to each representative sequence based on the silva138/16s_bacteria database with a confidence threshold of 70% using Naive bayes classifier in Qiime2.

### Data analysis

GraphPad Prism 9.3.1. was adopted to analyze the data, and the continuous data conforming to the normal distribution were described using mean and standard deviation. Differences between the intervention groups and the control group were analyzed by one-way ANOVA. A multivariate statistical analysis was performed using ROPLS in R package from Bioconductor on Majorbio Cloud Platform (https://cloud.majorbio.com). The Alpha diversity indices including observed species and the Shannon index were calculated with Mothur v1.30.1 (Schloss et al., 2009).

## Results

### Clinical signs

No mortalities were found in any of the control or lambda-cyhalothrin exposed groups during the 28 day treatment period, and no abnormal symptoms were registered in the first two weeks. From the third week onwards, the activity of all female mice in the treated groups was slightly reduced, and especially females of the high-dose group (2LCT) showed temporary clinical signs of semi closed eyelids (2 animals). In addition, aggressive performance was observed for the treated male mice in both exposed groups in the second half of the experiments (week 3 and 4). In the fourth week, male mice of the 2LCT group (one cage) started to bite and fight with their group mates, leading to serious injury on the back of one animal.

Body weight and food consumption data are shown in **Table 1**. Body weight and food consumption of treated animals showed a decrease compared to the controls during the experimental period, although the decrease in food consumption did not reach statistical significance except for the last determination for the female 2LCT group. The reduction of mouse body weight was most pronounced in females, with the most significant reduction in body weight presented at the second determination (day 13). Collectively, the administration of lambda-cyhalothrin markedly reduced the body weight of studied animals in a dose dependent way.

**Table 1.**
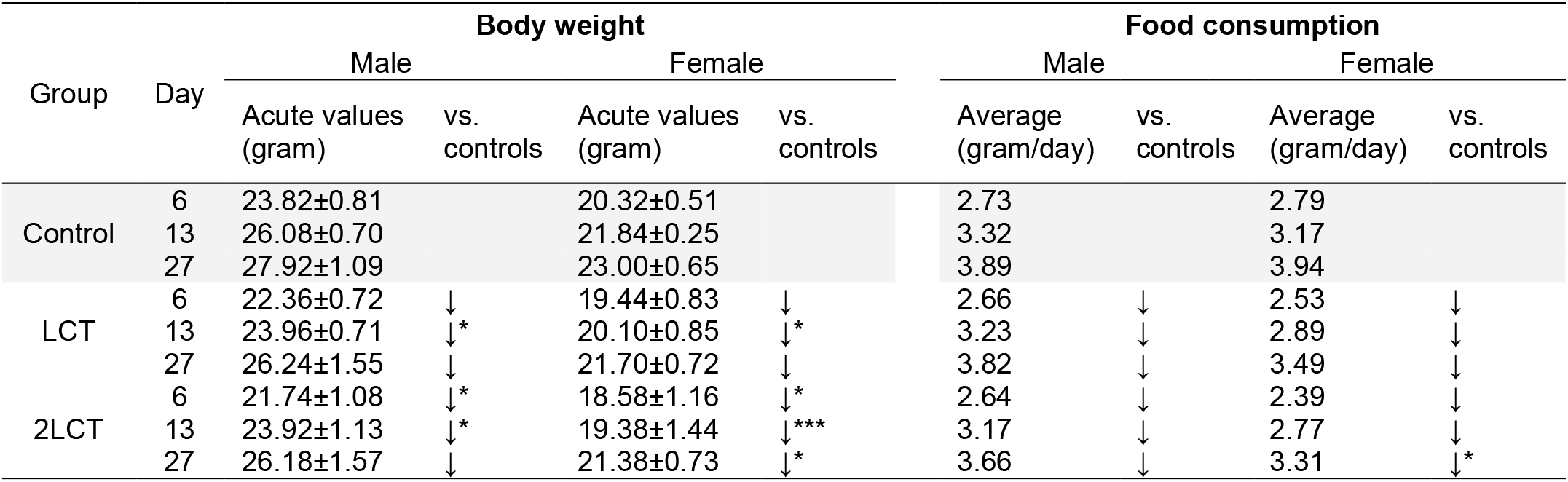
Body weight and food consumption of male and female C57BL/6 mice (n=5 per sex per group) dosed for 28 days with lambda-cyhalothrin or vehicle control. Data were collected on study days 6, 13, and 27 and values for exposed mice are compared to control (*p<0.05, **p<0.01, ***p<0.001)

| Group | Day | Body weight |  |  |  | Food consumption |  |  |  |
| --- | --- | --- | --- | --- | --- | --- | --- | --- | --- |
|  |  | Male |  | Female |  | Male |  | Female |  |
|  |  | Acute values (gram) | vs. controls | Acute values (gram) | vs. controls | Average (gram/day) | vs. controls | Average (gram/day) | vs. controls |
| Control | 6 | 23.82±0.81 |  | 20.32±0.51 |  | 2.73 |  | 2.79 |  |
|  | 13 | 26.08±0.70 |  | 21.84±0.25 |  | 3.32 |  | 3.17 |  |
|  | 27 | 27.92±1.09 |  | 23.00±0.65 |  | 3.89 |  | 3.94 |  |
| LCT | 6 | 22.36±0.72 | ↓ | 19.44±0.83 | ↓ | 2.66 | ↓ | 2.53 | ↓ |
|  | 13 | 23.96±0.71 | ↓* | 20.10±0.85 | ↓* | 3.23 | ↓ | 2.89 | ↓ |
|  | 27 | 26.24±1.55 | ↓ | 21.70±0.72 | ↓ | 3.82 | ↓ | 3.49 | ↓ |
| 2LCT | 6 | 21.74±1.08 | ↓* | 18.58±1.16 | ↓* | 2.64 | ↓ | 2.39 | ↓ |
|  | 13 | 23.92±1.13 | ↓* | 19.38±1.44 | ↓*** | 3.17 | ↓ | 2.77 | ↓ |
|  | 27 | 26.18±1.57 | ↓ | 21.38±0.73 | ↓* | 3.66 | ↓ | 3.31 | ↓* |

### Bile acid profiles in plasma

**Fig. 2** shows the results obtained for the bile acid profiles in plasma samples of control and lambda-cyhalothrin treated male and female mice. Of the twenty-four bile acids that could be detected by the method applied, in plasma samples only ten bile acids were present at levels that enabled their quantification. This included the conjugated bile acids TMCA (sum of TαMCA+TβMCA+TωMCA), TCA, TCDCA, THDCA, and TUDCA, the primary bile acids CA, αMCA, βMCA, and CDCA, as well as one secondary bile acid being DCA. Glycine-conjugated bile acids and other secondary bile acids were not detected or detected below the limitation of quantification (0.01 μM).

**Fig. 2.**
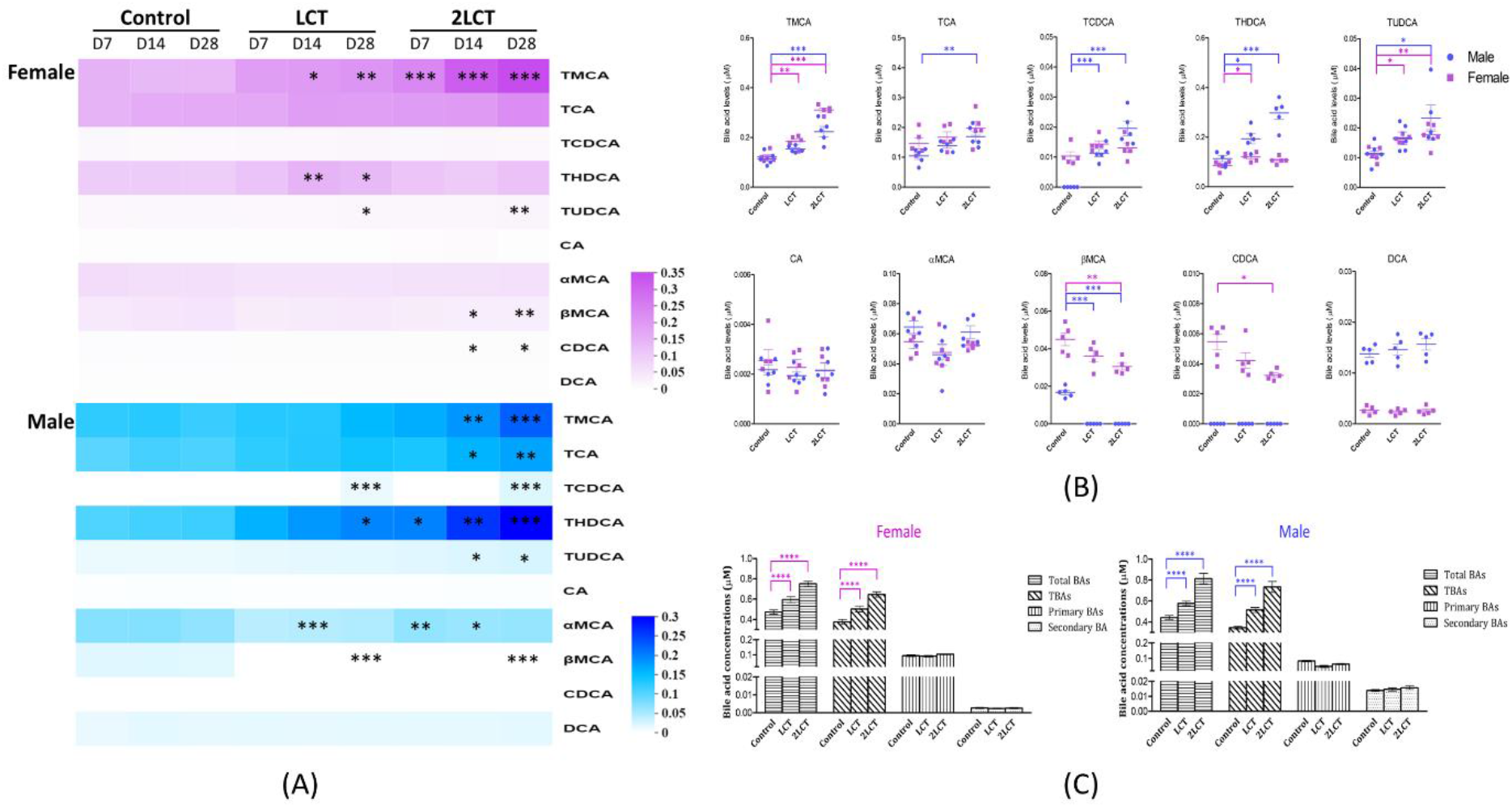
Bile acid profiles in plasma samples from male and female mice in untreated and lambda-cyhalothrin treated groups, shown as (A) a heatmap presenting mean concentrations of the detected bile acids in plasma samples from studied mice on days 7, 14, and 28 with shades of pink (female) and blue (male) indicating the concentrations; (B) the concentrations of individual bile acids detected in plasma samples on day 28, and (C) the total amount of bile acids (total BAs) in plasma samples at day 28 including also the total concentration of taurine conjugated bile acids (TBAs), primary bile acids (primary BAs), and secondary bile acids (secondary BAs) (n=5; *p<0.05, **p<0.01, ***p<0.001, ****p<0.0001). The low and high dose groups are presented by LCT and 2LCT, respectively.

**Fig. 2A** shows in a heatmap the time-dependent mean concentrations of the bile acids detected in plasma samples from male and female mice of all groups on days of 7, 14, and 28. The most significant changes were detected in taurine conjugated bile acids that showed significant increases in a dose dependent manner in all plasma samples of treated mice compared to controls. Changes in the concentrations of primary and secondary bile acids were limited to significant decreases in βMCA in male and female mice exposed to lambda-cyhalothrin, and in CDCA for exposed female mice. The effects observed were dose and also time dependent with the largest changes detected in the 2LCT groups at the last experimental day (day 28). In addition, some substantial gender dependent differences were observed (**Fig. 2B**) especially for i) THDCA which increased more in male than female mice exposed to lambda cyhalothrin, ii) βMCA and CDCA which were present at lower levels and often below the limit of quantification in all male as compared to female plasma samples and iii) DCA which was present at substantially higher concentrations in plasma samples of all female as compared to male animals. Collectively, significant dose dependent increases in all detected conjugated bile acids were observed in plasma samples from male and female lambda-cyhalothrin treated mice compared to controls (**Fig. 2A** and **2B**). Of the primary and secondary bile acids, dose dependent decreases were observed for βMCA and CDCA whereas for CA, αMCA and DCA there were no differences between the control and lambda-cyhalothrin exposed groups (**Fig. 2A** and **2B**). **Fig. 2C** summarizes the bile acid profiles for the different groups at day 28 focussing on the total amount of bile acids as well as the total amount of taurine conjugated, primary, and secondary bile acids in the plasma of the mice. From this analysis it follows that there is a dose dependent increase in the plasma concentration of the total bile acids which can be mainly ascribed to a dose dependent increase in the amount of taurine conjugated bile acids.

### Bile acid profiles in feces

In addition to plasma bile acid concentrations, bile acid profiles were also quantified in fecal samples. **Fig. 3** shows the results obtained. Fourteen bile acids including six taurine conjugated bile acids including TCA, TMCA (sum of TαMCA+TβMCA+TωMCA), TDCA, THDCA, TLCA, and TUDCA, three primary bile acids including CA, αMCA, and βMCA, and five secondary bile acids including DCA, LCA, UDCA, ωMCA, and HDCA could be quantified in the fecal samples from male and female mice of lambda-cyhalothrin treated and control groups collected on day 28.

**Fig. 3.**
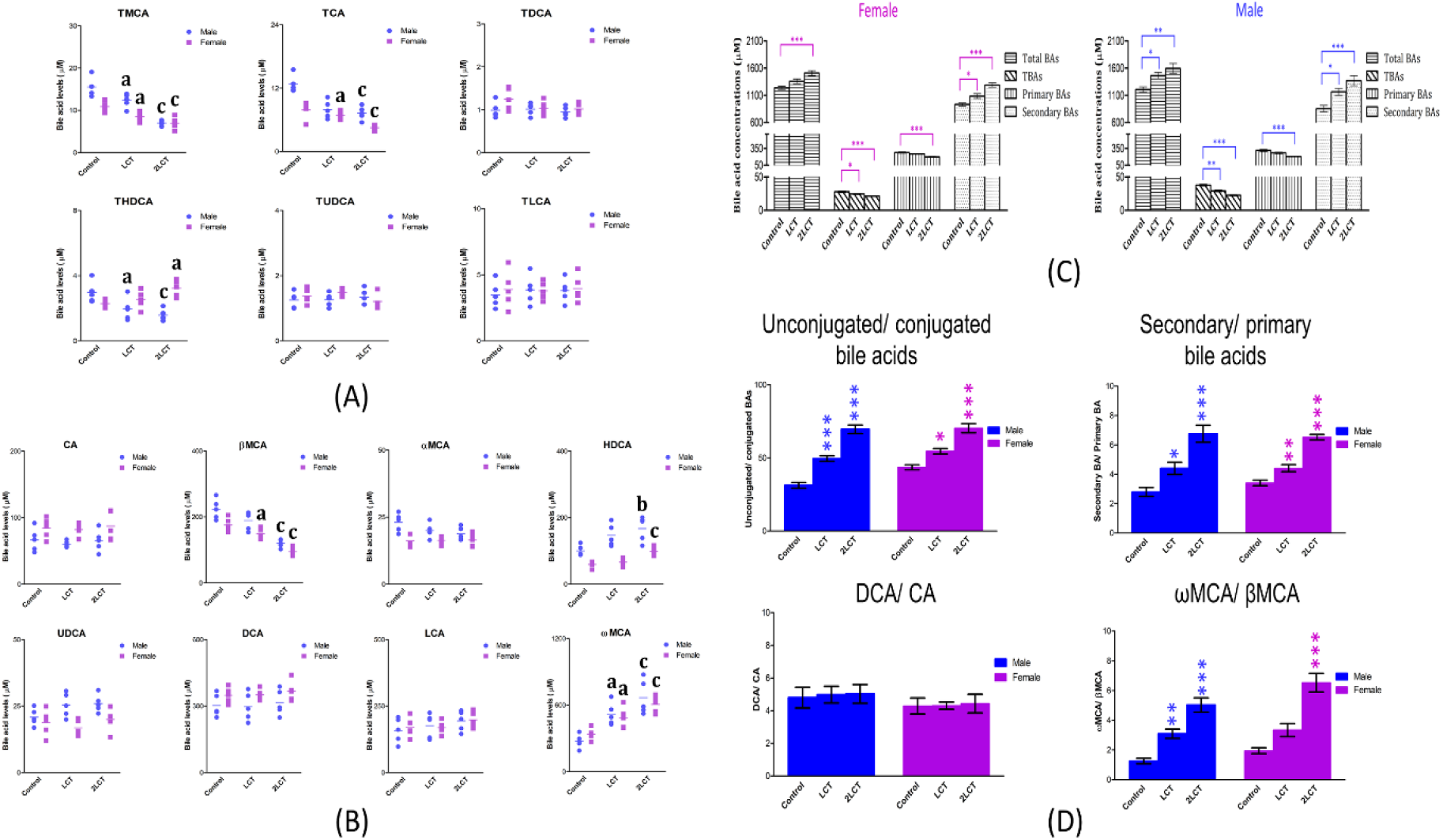
Bile acid profiles in fecal samples from male and female mice in untreated and lambda-cyhalothrin treated groups on day 28 as characterized by (A) concentrations of fecal taurine conjugated bile acids; (B) fecal unconjugated bile acids; (C) the total amount of fecal bile acids (total BAs), fecal taurine conjugated bile acids (TBAs), fecal primary bile acids (primary BAs), and fecal secondary bile acids (secondary BAs); and (D) the ratio of unconjugated to conjugated bile acids (unconjugated/ conjugated bile acids), the ratio of primary to secondary bile acids (secondary/ primary bile acids), the ratio of DCA to CA (DCA/ CA), and the ratio of ωMCA to βMCA (ωMCA/ βMCA). Data for male (blue) and female (pink) mice fecal samples are presented and significance of the differences in treated animals compared to controls is also indicated (a, p<0.05; b, p<0.01; c, p<0.001 in A and B; *, p<0.05; **, p<0.01; ***, p<0.001 in C and D). The low and high dose groups are presented by LCT and 2LCT, respectively.

The levels of fecal taurine conjugated bile acids and fecal unconjugated bile acids in samples from the studied male and female mice are presented in **Fig. 3A** and **3B**. For taurine conjugated bile acids (**Fig. 3A**), significant dose dependent decreases were observed in TMCA and TCA in lambda-cyhalothrin treated male and female animals, whereas levels of THDCA showed substantial gender dependent differences with a significant decrease in feces of treated male mice and an increase in fecal samples of treated females, especially in the 2LCT group. For unconjugated bile acids (**Fig. 3B**), significant dose dependent alterations were detected in lambda-cyhalothrin treated animals, with a marked decrease of the primary bile acid βMCA and an accompanying increase of the secondary bile acids ωMCA and HDCA. **Fig. 3C** summarizes the fecal bile acid profiles of lambda-cyhalothrin treated animals and controls at day 28 based on the amount of total, taurine conjugated, primary and secondary bile acids. A significant dose dependent increase in the level of fecal secondary bile acids at the expense of fecal taurine conjugated and primary bile acids was observed, leading to a markedly increased size of the fecal bile acid pool. It has been reported that the majority of secondary bile acids in mice is composed of ωMCA and DCA that are formed by respectively 6β-epimerization of βMCA and 7α-hydroxylation of CA (Sayin et al., 2013). The ratio of βMCA over ωMCA and CA over DCA, as well as that of the total fecal primary bile acids over fecal secondary bile acids, together with another important ratio being the ratio between total fecal unconjugated bile acids and fecal conjugated bile acids (Sayin et al., 2013) are presented in **Fig. 3D**. A markedly higher conversion rate from primary bile acids to secondary bile acids was observed in fecal samples from lambda-cyhalothrin treated animals, which could be ascribed to the significantly faster 6β-epimerization of βMCA to ωMCA in these treated animals than in control mice. The rate of deconjugation was also obviously raised in the treated animals, whereas the transformation from CA to DCA did not show differences between lambda-cyhalothrin treated animals and controls.

### Bacterial profiling

**Fig. 4** describes the bacterial profiling of the fecal samples from lambda-cyhalothrin exposed mice and controls at the phylum and family level, as well as an analysis of alpha diversity performed based on the determined operational taxonomic units (OTUs). The results obtained reveal that the phyla of *Bacteroidota* and *Firmicutes* were dominant in fecal samples of the studied mice and no significant changes in the microbial community at phylum level were observed in lambda-cyhalothrin treated mice compared to controls. Also the alpha diversity indicated by the index of total species observed (sobs) as well as the shannon diversity showed that no significant changes were found in the total amount or richness of the microbial community after the administration of lambda-cyhalothrin (**Fig. 4B**). At the family level, the proportions of the main twenty-one bacterial families in the fecal samples from male and female mice are shown in **Fig. S1**, and following from these results **Fig. 4C** presents the proportions of the ten most common families of gut microbial species in male and female mice.

**Fig. 4.**
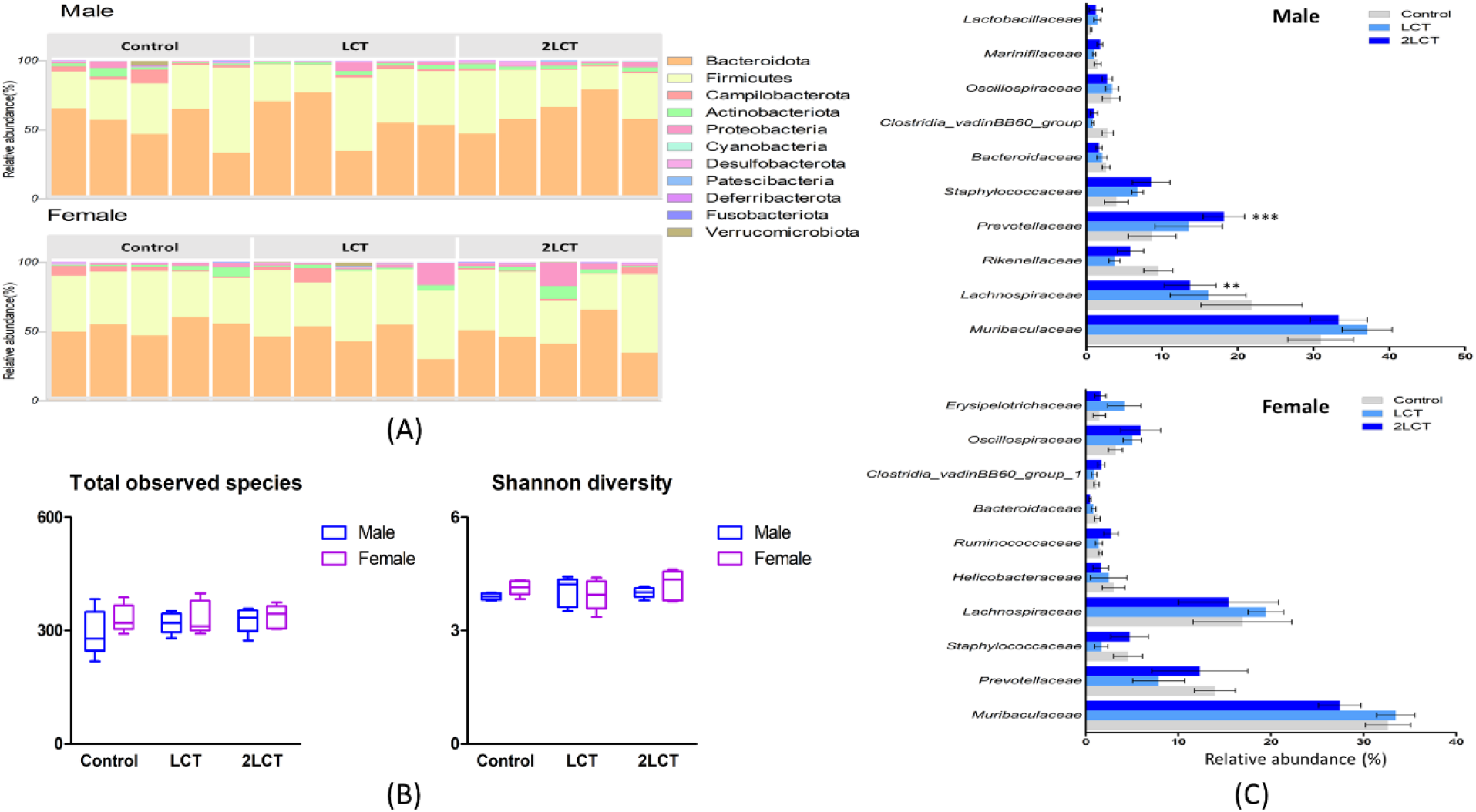
Gut bacterial profiles of fecal samples from untreated (control) and lambda-cyhalothrin treated male and female mice on day 28. LCT and 2LCT are the low and high dose groups. (A) shows the relative abundance of the gut microbial community in mice fecal samples at the phylum level (n=5 per sex per group), and (C) at the family level, presenting only the ten dominant families. (B) shows the alpha diversity by values of total observed species and shannon diversity (*p<0.05, **p<0.01, ***p<0.001).

As shown in **Fig. 4C**, a significant dose dependent increase in relative abundance of *Prevotellacea* (phyla of *Bacteroidota*) and a dose dependent decrease in relative abundance of *Lachnospiraceae* (phyla of *Firmicutes*) were observed in male mice treated with lambda-cyhalothrin which were significant compared to control at the high dose level (2LCT group). In contrast, no significant bacterial changes at family level were observed in the lambda-cyhalothrin treated females.

### Correlations between lambda-cyhalothrin induced changes in microbial and bile acid profiles

The changes in the gut microbial community upon exposure to lambda-cyhalothrin were also analyzed at genus level, and relationships between the changes in the bacterial and bile acid profile were analysed by a correlation analysis. Results obtained are shown in **Fig. 5**. The bacterial species which were substantially different upon treatments were selected for this correlation analysis and are summarized in **Fig. 5A**. The relative abundance of the genera named *Oscillospiraceae NK4A214 group* (class, Clostridia; phylum, Firmicutes) significantly increased in fecal samples from lambda-cyhalothrin treated male mice of the LCT and 2LCT groups, followed by the genera *Brevibacterium* and *Brachybacterium* that were increased albeit not significant (p>0.05). In treated females, the relative abundance of the genera *Brachybacterium*and and *Brevibacterium* belonging to the phyla of *Actinobacteriota* was also increased, with statistical significance (p<0.05) obtained for *Brachybacterium* in the female 2LCT group. **Fig. 5B** presents the correlations and the statistical significance between the fifty dominant genera in mice feces and the significantly altered fecal bile acid related endpoints including the ratios of total unconjugated bile acids and conjugated bile acids, total secondary and primary bile acids, as well as ωMCA and βMCA, and the significantly increased levels of HDCA (see **Fig. 3**). In the heatmap presentation (**Fig. 5B**), it can be seen that the genera of *Brachybacterium* and *Brevibacterium* showed a positive correlation with the ratio of unconjugated bile acids to conjugated bile acids (p<0.01), revealing their relationship with bile acid deconjugation. Meanwhile, *Brachybacterium* and *Brevibacterium* were shown to be positively associated with the ratio of secondary and primary bile acids (p<0.05) as well as the ratio between the secondary bile acid ωMCA and the primary bile acid βMCA (p<0.001). The latter indicates the correlations of the raised richness of *Brachybacterium* and *Brevibacterium* with the markedly faster 6β-epimerization from βMCA to ωMCA leading to the significantly increased conversion of the total fecal primary bile acids to secondary bile acids. In contrast, the genera *Oscillospiraceae NK4A214 group* appeared to be only ones highly linked to the production of HDCA, the 7β-hydroxylated metabolite of ωMCA (Sayin et al., 2013), as reflected by its significantly positive correlation with the 7β-hydroxylation of ωMCA. Besides, the genera *Prevotellceae UCG spp* (class, Bacteroidia; phylum, Bacteroidota) also showed a significant positive correlation with the deconjugation and the production of secondary bile acids, particularly ωMCA produced from βMCA (p<0.01). Further, a linear regression model was utilized to study the correlation of the mouse fecal bile acid profiles with the dominant bacterial genera that showed significant changes in **Fig 5B**, and the analyzed correlations with R square values above 0.35 are summarized in **Fig. 5C**. Results obtained corroborate the positive link of *Brachybacterium* as well as *Brevibacterium* with the 6β-epimerization of βMCA, and the correlation of the raised abundance of *Oscillospiraceae NK4A214 group* with the faster 7β-hydroxylation of ωMCA in males. Moreover, the correlations between the concentrations of individual fecal bile acids and the relative abundance of fifty bacterial species are shown in **Fig. S2**. In addition to the genera referred to above, *Bacteroides* (phylum Bacteroidota), *Staphylococcus* (phylum Firmicutes) and *Bacilli RF39 spp* (phylum Firmicutes) followed by *Rumminococcaceae spp, Oligella, Helicobacter* showed links to the transformation of individual fecal bile acids as well.

**Fig. 5.**
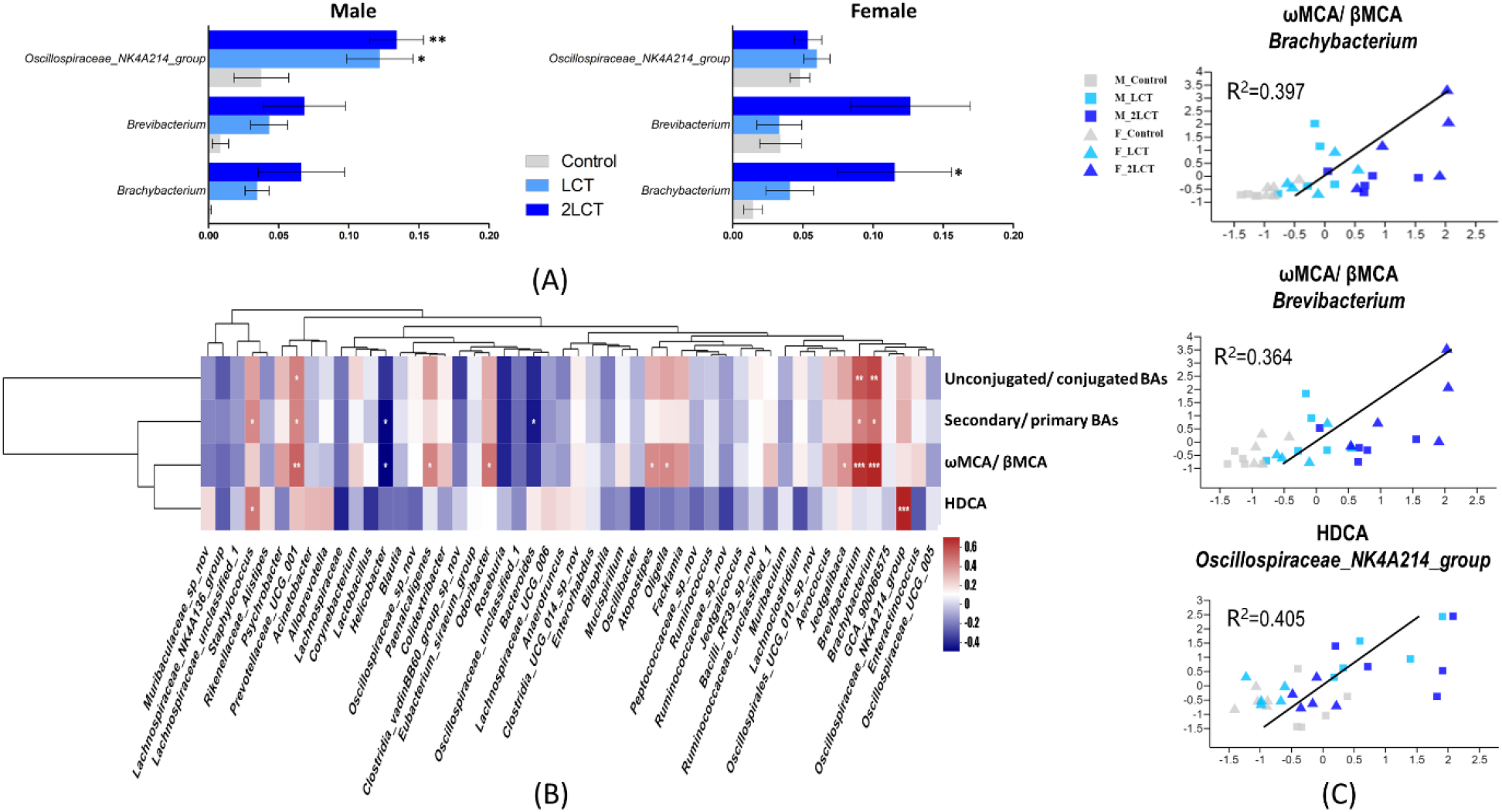
Gut bacterial genera in fecal samples of male and female control and lambda-cyhalothrin treated mice on day 28, as well as correlations between the bacterial and bile acid profiles. LCT and 2LCT are the low and high dose groups. (A) shows the significantly altered bacterial genera in lambda-cyhalothrin treated male and female mice of LCT (light blue), and 2LCT (blue) groups compared to controls (grey). (B) shows the heatmap presentation with pearson correlation coefficients of the main fifty genera in mouse fecal samples with the values of unconjugated/ conjugated bile acids, secondary/ primary bile acids, ωMCA/ βMCA, and the significant increased concentration of fecal HDCA. The positive and negative correlation are separately presented by colours of red and blue (*p<0.05, **p<0.01, ***p<0.001). The genera significantly related to conversion of fecal bile acids (in B) were further analyzed using linear regression, and the correlations with R squares >0.3 between bacterial and bile acid profiles are shown in (C). The untreated and treated male (M) and female (F) mice of different groups were shown, namely, M_Control, M_LCT, M_2LCT, F_Control, F_LCT, and F_2LCT respectively (n=5).

### Additional analysis of the impact of lambda-cyhalothrin on gut bacterial and bile acid profiles

**Fig. 6A** and **B** separately show the results of a principal component analysis (PCA) for the bile acid profiles in respectively plasma and fecal samples from control and lambda-cyhalothrin treated mice. As shown in the two graphs, the bile acid profiles of untreated male and female mice cluster together and show a dose dependent separation from those of lambda-cyhalothrin treated mice, with the low-dose LCT group and high-dose 2LCT group clustering per group with a separation in between. The analysis also reveals the gender dependence of the effects as well as a more pronounced separation for the plasma than the fecal bile acid profiles. **Fig. 6C** describes the effects of lambda-cyhalothrin on the gut microbial community based on data of OTUs in mice fecal samples, and shows no clear cluster of one gender or one group (treated or untreated) which is in line with the effects at phylum level and indicates the huge intra-species variations of bacterial profiles as compared to the detected lambda-cyhalothrin induced effects using data on OTUs. Moreover, **Fig. 6D** describes the correlations between the bile acid profiles in plasma and fecal samples by groups of taurine conjugated, primary and secondary bile acids. The levels of taurine conjugated bile acids in plasma showed highly positive correlation with those of fecal secondary bile acids, and was negatively associated with the concentrations of fecal taurine conjugated and primary bile acids.

**Fig. 6.**
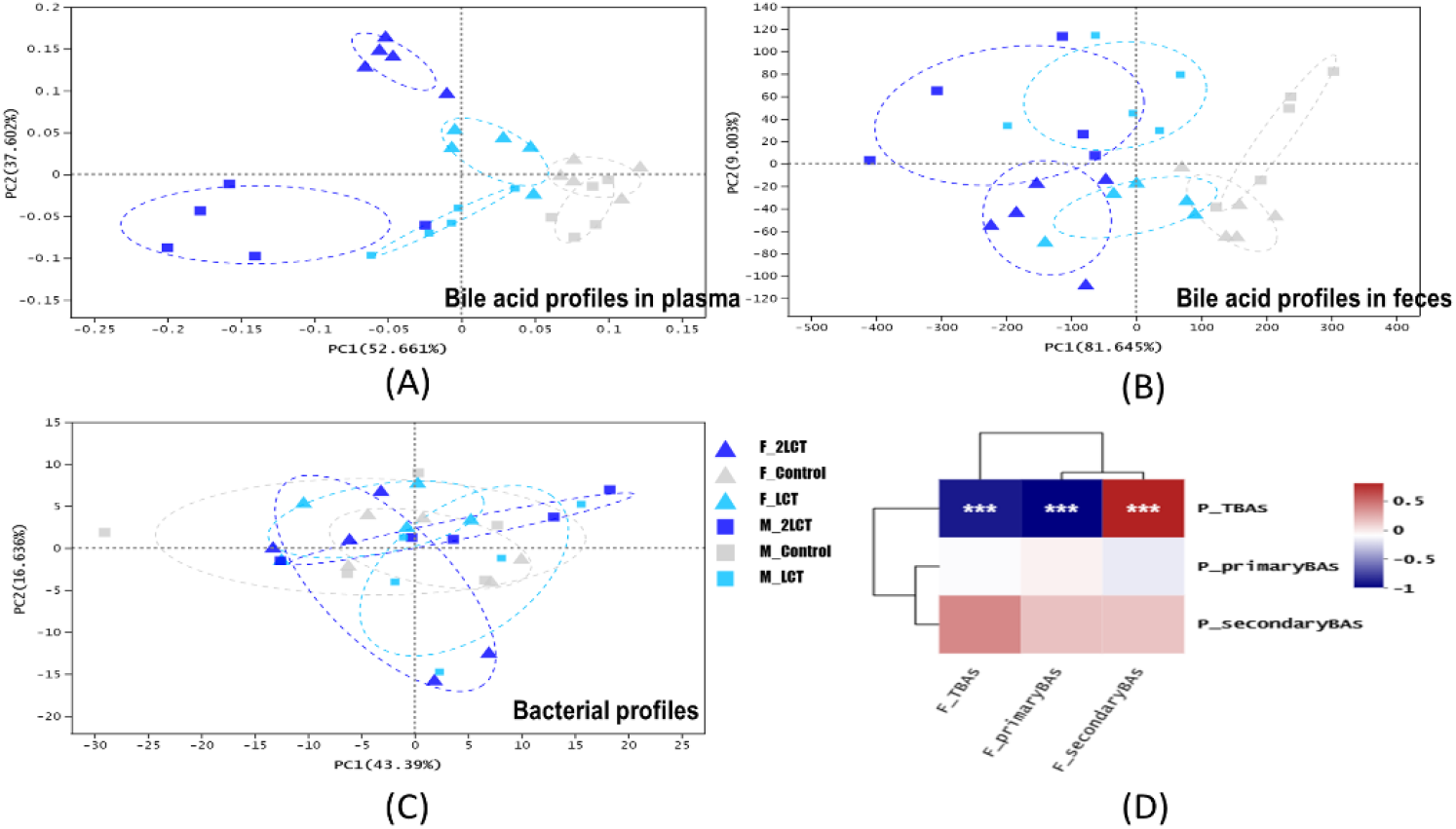
Principal component analysis (PCA) of bile acid and bacterial profiles, as well as the correlation between bile acid profiles in plasma and fecal samples. (A) and (B) show the PCA analyses for respectively the plasma and fecal bile acid profiles, and (C) shows the PCA analysis using OTUs for the fecal bacterial profiles. All plasma and fecal samples were obtained from control (grey) and lambda-cyhalothrin treated (LCT, low dose, light blue; 2LCT, high dose, blue) male (squares) and female (triangles) animals on day 28 (n=5 mice per group and sex). In (D), the correlation between bile acid profiles of plasma (P) and fecal (F) samples from all studied mice on day 28 was analyzed by pearson correlation coefficients, and is shown for groups of total taurine conjugated bile acids, total primary bile acids, and total secondary bile acids.

## Discussion

In this study, the alterations in plasma and fecal bile acid profiles as well as in the gut microbial community induced by the exposure of male and female mice to lambda-cyhalothrin were characterized, showing significant lambda-cyhalothrin induced effects compared to controls (**Fig. 6A, B**, and **C**).

A key finding of the current study is that the fecal bile acid profile was markedly altered upon the treatment of the mice with lambda-cyhalothrin, comprising not only a dose-dependent increase in the amount of total and secondary bile acids and decreased levels of conjugated and primary bile acids (**Fig. 3C**), but also dose-dependently raised ratios of unconjugated/ conjugated bile acids and secondary/ primary bile acids. Notably, the raised transformation from βMCA to ωMCA relating to an accelerated 6β-epimerization (Thomas et al, 2008) was also found in our previous study with an optimized 24 h in vitro fermentation model in which mice feces was incubated with cyhalothrin or another commonly used pyrethroid cypermethrin, leading also to an increased ratio of secondary/ primary bile acids (Zheng et al. submitted). In addition to changes in βMCA and ωMCA, the in vitro study also showed a significant decrease in αMCA upon exposure of the gut microbiota to cyhalothrin, while this was not shown in the present study. The reason for this discrepancy could be that in the current in vivo work the deconjugation was increased, as indicated by the raised ratio of unconjugated/ conjugated bile acids. Similar to the in vitro result, αMCA appeared to be efficiently metabolized, for example by epimerization to βMCA (de Boer et al., 2020), which, in combination with the faster deconjugation from TαMCA to αMCA could maintain the dynamic balance for the levels of αMCA in this in vivo study.

Although only approximately 5 % of bile acids secreted by hepatocytes end up in feces (Hundt et al., 2017), the fecal bile acid profile was previously reported to largely reflect the intestinal bile acid composition which is dependent on activities of microbial enzymes (Joyce and Gahan et al., 2016) such as BSH, 7α-dehydroxylase, and 6β-epimerase present in the gut microbiota. The accelerated deconjugation related to the raised activities of microbial BSH (Sayin et al., 2013) could be linked to the significantly increased richness of the genera *Brachybacterium* and *Brevibacterium* upon the high-dose lambda-cyhalothrin treatment of female animals (**Fig. 5A** and **B**). Meanwhile, the conversion of βMCA to ωMCA is carried out by 6β-epimerase (Chiang and Ferrell et al., 2020), which could also be provided by the genera *Brachybacterium* with *Brevibacterium* in the current study (**Fig. 5B**). Conversely upon the administration of lambda-cyhalothrin, there were no obvious effects on 7α-dehydroxylation of CA to DCA, which indicates that the community of bacterial strains carrying 7α-dehydroxylase was not markedly affected. In addition to these microbial enzymes, the increase in fecal HDCA, the 7β-hydroxylated metabolite of ωMCA could be ascribed to the highly active 7β-hydroxylase (Sayin et al., 2013) provided by the enriched genera *Oscillospiraceae NK4A214 group* upon the treatment of male mice with lambda-cyhalothrin. Moreover, interactions between bile acids and intestinal microbiota have been previously described (Ramírez-Pérez et al., 2018), and the impact of bile acid composition in the gut on the intestinal microbiome has been examined. For example i) conjugated bile acids were reported to exhibit antibacterial activity (Jones et al., 2008; Inagaki et al., 2006); ii) the ratio of secondary over primary bile acids was reported to be positively correlated to certain commensal genera (Kakiyama et al., 2013). Thus, the altered gut microbial community acquired upon the treatment with lambda-cyhalothrin could be caused by the changed fecal bile acid composition as well.

In addition to fecal bile acids, the bile acid profiles in plasma also significantly altered especially with respect to taurine conjugated bile acids (**Fig. 3** and **6A**). These levels of plasma taurine conjugated bile acids were positively correlated to the proportions of fecal secondary bile acids (**Fig. 6D**). The raised amount of intestinal secondary bile acids are expected to be reabsorbed into enterocytes, secreted into the portal circulation, and delivered to hepatocytes (Thomas et al., 2008) in which the secondary bile acids are conjugated with taurine by specific enzymes to generate conjugated secondary bile acids leading to for example the increased levels of THDCA, TUDCA, and TωMCA (included in TMCA) detected in plasma (**Fig. 2A** and **B**). Additionally, the significantly decreased levels of the fecal primary bile acid βMCA could directly lead to its reduction in plasma upon the treatment with lambda-cyhalothrin especially in male mice. However, the taurine conjugated primary bile acids such as TCA, were decreased in fecal samples but simultaneously increased in plasma, indicating that these effects may rather be ascribed to changes in bile acid transporters.

The enterohepatic circulation of bile acids is presented in **Fig. 7** showing the flow of bile acids through the gut-liver axis and the transporters involved, while also presenting the changes of intestinal and plasma bile acid profiles induced by the treatment of the mice with lambda-cyhalothrin. As shown, except for the intestinal bacterial community, the hepatocytes and intestinal epithelial cells also contribute to the bile acid metabolism and transport, with an important role for the organic solute transporter subunit α/β (OSTα/ β). The crucial role of OSTα/ β in intestinal bile acid transport is well known, and its expression pattern highly correlates with that of the intestinal bile acid transporter ASBT (apical sodium-dependent bile acid transporter); additionally, the OSTα/ β is capable to protect the enterocytes from bile acid accumulation (Ferrebee et al., 2018). Previously, it has been shown that the expression of OSTα/ β could be not only affected by exposure to xenobiotics (Beaudoin et al., 2020), but also regulated by the bile acids such as CA or CDCA themselves (Frankenberg et al., 2006). As a result, it can be proposed that the raised proportions of taurine conjugated bile acids in plasma accompanied by reduced levels in feces could be caused by an increase in the expression and activity of ASBT and hepatic/ intestinal OSTα/β, leading to more efficient reuptake of the conjugated bile acids from the intestinal tract into the portal vein and liver. In the last two decades, the role of OSTα/ β and ABST for the disposition of endogenous compounds particularly relating to the enterohepatic circulation of bile acids gained increasingly interest (Beaudoin et al., 2020; Vicens et al., 2007). Thus, such lambda-cyhalothrin induced effects on the expression and activity levels of ASBT and/or OSTα/β presents a topic of interest for future studies.

**Fig. 7.**
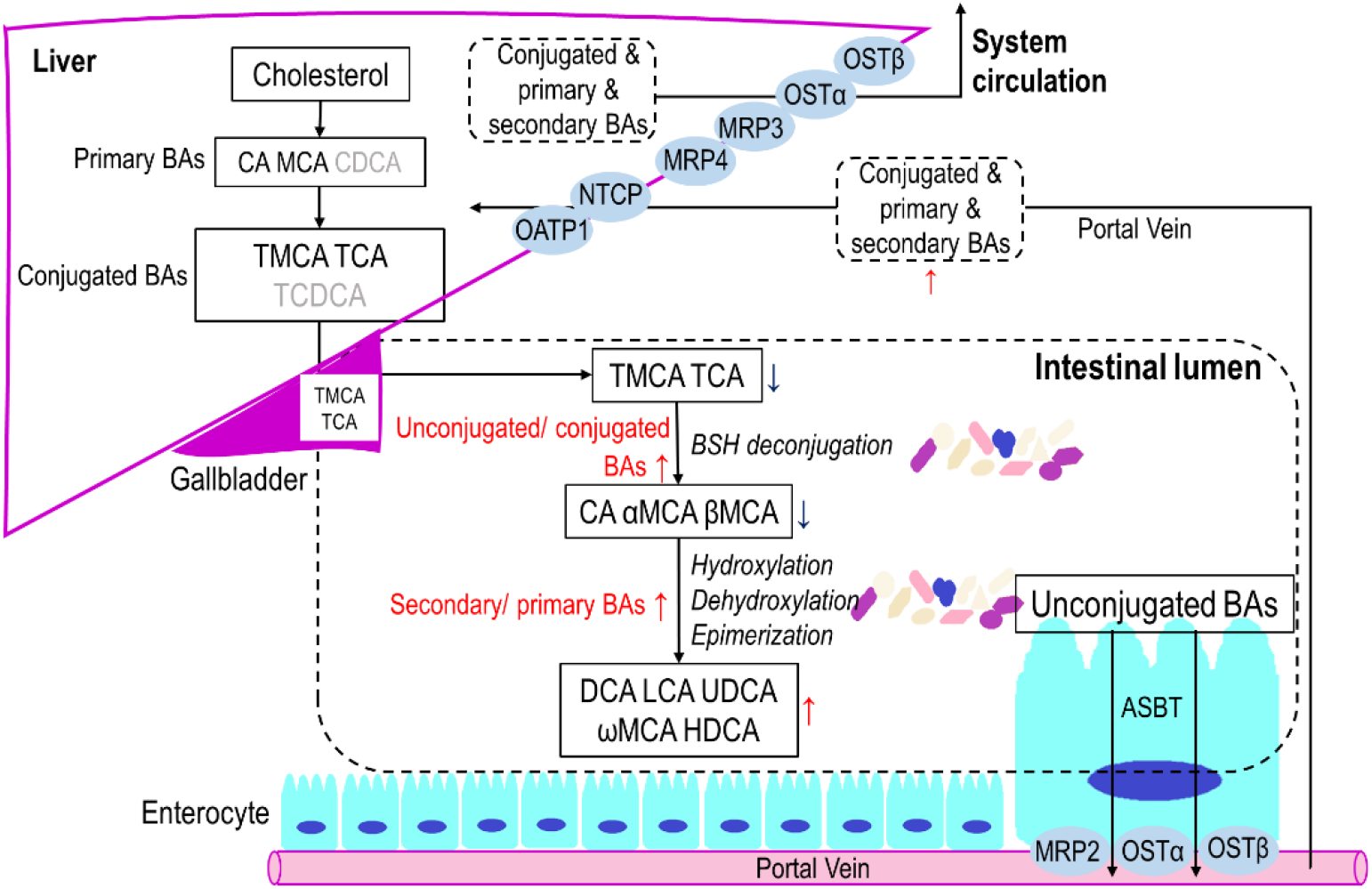
The biosynthesis, transport and metabolism of bile acids, as well as the alterations of bile acids in plasma and fecal samples of mice upon 28 days administration of lambda-cyhalothrin as reported in the present study. The upregulated and downregulated levels of bile acids were separately marked red and blue. Bile acids are produced by the liver, mainly conjugated to taurine (TMCA and TCA), excreted into the bile, released by the gallbladder into the intestine of mice via the bile duct. In the intestinal lumen, TMCA and TCA are deconjugated to CA, αMCA, and βMCA via BSH, and hydroxylated or dehydroxylated or epimerized to form the secondary bile acids of DCA, LCA, UDCA, ωMCA, and HDCA by mouse gut microbiota. At the terminal ileum, most unconjugated bile acids are reabsorbed by the transporter ASBT into enterocytes and secreted into the portal circulation via the basolateral bile acid transporters OSTα, OSTβ, and MRP2 (Trauner and Boyer, 2003). Bile acids are then taken up by NTCP and OATP1 into hepatocytes. Hepatic MRP3, MRP4 and the OSTα–OSTβ complex provide alternative excretion routes for bile acids into the systemic circulation (Meier and Stieger, 2002).

Furthermore, considering the important role of bile acid homeostasis in host health, it is of interest to note that increased levels of secondary bile acids as induced by the treatment of the mice with lambda-cyhalothrin in the present study, have been previously noted in the stool of hosts with ulcerative colitis and dysplasia or cancer (Bernstein et al., 2011; Ajouz et al., 2014); additionally, the secondary bile acids can also stimulate the bile acid receptor TGR5, which may result in suppression of the inflammatory response to microbial ligands, potentially leading to a lower degree of immunosurveillance against neoplasms (Vaughn et al., 2019). Besides, if our hypothesis for the increase in ASBT and/or hepatic/ intestinal OSTα/β holds, it can result in cholestatic disorders and other metabolic diseases posing a risk to host health as well (Beaudoin et al., 2020). Therefore, further studies on xenobiotic-gut microbiota-bile acid metabolism interactions and their consequences for host health are of interest for future studies.

## Conclusion

In this study, the effects of lambda-cyhalothrin on gut bacterial and bile acid profiles were assessed in male and female mice. The key findings were that the treatment of the mice with lambda-cyhalothrin induced a markedly raised total amount of plasma and fecal bile acids, mainly caused by an increase in plasma conjugated bile acids and in fecal secondary bile acids. In addition, the gut microbial community showed a significantly increased richness of *Prevotellacea* and a depletion of *Lachnospiraceae* at the family level upon the treatment with lambda-cyhalothrin, and the genera *Brachybacterium, Brevibacterium* as well as *Oscillospiraceae NK4A214* were found to be highly correlated to the fecal bile acid levels. In summary, the alterations of bacterial and bile acid profiles obtained provide a proof-of-principle for effects of lambda-cyhalothrin on gut microbiota and related bile acid metabolism which may eventually influence host health.

## Acknowledgements

The authors acknowledge prof. dr Jacques Vervoort for his supervision and efforts during the whole work. We are grateful to Wouter Bakker for help with the LC-MS/MS, and to staff members working in the animal laboratory of BIT (Beijing Institute of Technology, China) for help with animal handling.

We thank the fundings supported by Key Lab of Agro-product Quality and Safety (Beijing, China).

## Abbreviations

Bile acids:

(CA): cholic acid
(TCA): taurocholic acid
(GCA): glycocholic acid
(CDCA): chenodeoxycholic acid
(TCDCA): taurochenodeoxycholic acid
(GCDCA): glycochenodeoxycholic acid
(DCA): deoxycholic acid
(TDCA): taurodeoxycholic acid
(GDCA): glycodeoxycholic acid
(LCA): lithocholic acid
(TLCA): taurolithocholic acid
(GLCA): glycolithocholic acid
(UDCA): ursodeoxycholic acid
(TUDCA): tauroursodeoxycholic acid
(GUDCA): glycoursodeoxycholic acid
(HDCA): hyodeoxycholic acid
(THDCA): taurohyodeoxycholic acid
(GHDCA): glycohyodeoxycholic acid
(αMCA): α-muricholate
(TαMCA): tauro-α-muricholate
(βMCA): β-muricholate
(TβMCA): tauro-β-muricholate
(ωMCA): ω-muricholate
(TωMCA): tauro-ω-muricholate

Others:

(LCT): lambda-cyhalothrin
(BSH): bile salt hydrolase
(DMSO): dimethyl sulfoxide
(LCMS): electrospray liquid chromatography mass spectrometry
(ESI): electrospray ionization
(OTUs): operational taxonomic units
(ASBT): apical sodium-dependent bile acid transporter
(OSTα/β): organic solute transporter subunit α/β
(MRP): multidrug resistance-associated protein
(NTCP): sodium taurocholate cotransporting polypeptide
(OATP1): organic anion-transporting polypeptide 1

